# DHODH is an independent prognostic marker and potent therapeutic target in neuroblastoma

**DOI:** 10.1101/2021.09.25.461496

**Authors:** Thale Kristin Olsen, Cecilia Dyberg, Bethel Embaie, Adele Alchahin, Jelena Milosevic, Jörg Otte, Conny Tümmler, Ida Hed Myrberg, Ellen M. Westerhout, Jan Koster, Rogier Versteeg, Per Kogner, John Inge Johnsen, David B. Sykes, Ninib Baryawno

## Abstract

Despite intensive therapy, children with high-risk neuroblastoma are at risk of treatment failure. We applied a pan-cancer, multi-omic system approach to evaluate metabolic vulnerabilities in human neuroblastoma. By combining metabolomics, CRISPR screen and gene expression data from more than 700 solid tumor cell lines, we identified *DHODH*, a critical enzyme in pyrimidine synthesis, as a potential novel treatment target in neuroblastoma. Of note, DHODH inhibition is currently under clinical investigation in patients with hematologic malignancies. In neuroblastoma, *DHODH* expression was identified as an independent risk factor for aggressive disease, and high *DHODH* levels correlated to worse overall and event-free survival. A subset of high-risk neuroblastoma tumors with the highest *DHODH* expression was associated with a dismal prognosis, with a 5-year survival of less than 10%. In neuroblastoma cell lines, *DHODH* gene dependency was found to correlate with *MYCN* dependency, rendering these cell lines highly sensitive to DHODH inhibition *in vitro*. In xenograft and transgenic neuroblastoma mouse models, tumor growth was dramatically reduced, and survival extended following treatment with the DHODH inhibitor brequinar. A combination of brequinar and temozolomide cured the majority of transgenic TH-MYCN neuroblastoma mice, indicating a highly active clinical combination therapy with curative potential. Overall, DHODH inhibition combined with temozolomde has clear therapeutic potential in neuroblastoma and we propose this combination as a candidate for clinical testing.

## INTRODUCTION

Neuroblastoma is an embryonal childhood tumor that arises in the tissue of the adrenal medulla, paraspinal or other sympathetic ganglia and accounts for about 6% of all childhood cancers^1–3^. Despite intensive multimodal therapy, survival in the high-risk group of neuroblastoma patients remains less than 50%. The *MYCN* oncogene is frequently amplified in neuroblastoma (around 20% of cases) and is one of the defining hallmarks of high-risk disease^1^. N-myc proto-oncogene protein (MYCN) is a transcription factor whose target genes are widely involved in metabolism, apoptosis and cell growth^4–6^ and its dysregulation in neuroblastoma leads to increased proliferation, decreased apoptosis and differentiation arrest^7,8^. *MYCN* amplification characterizes a subset of very aggressive tumors and correlates with poor prognosis. Moreover, the closely related c-myc transcription factor (*MYC*) has been established as a potent oncogene in an additional subset of neuroblastomas (approximately 10%)^9^. Furthermore, oncogenic *MYC* is known to orchestrate a profound rewiring of the metabolic processes in cancer cells. By driving the expression of metabolic genes, *MYC* increases production of ATP and other cellular key building blocks, which enable uncontrolled cell proliferation^10,11^. *MYCN* is also known to induce several metabolic alterations, such as increased glycolysis, oxidative phosphorylation, and glutamine metabolism^12–14^. Unconstrained growth in *MYC*- and *MYCN*-driven cancers is thus dependent on metabolic pathways, which may serve as novel targets for cancer therapy.

Genome-wide epigenetic and transcriptomic studies in neuroblastoma cell lines have demonstrated that there are at least two different malignant neuroblastoma cell types, namely the mesenchymal (MES) and adrenergic (ADRN) cells that are to a large extent controlled by networks of super-enhancers^15,16^. In a recent study of the super-enhancer landscapes in primary neuroblastoma patient samples, the presence of these two states were confirmed, and the ADRN phenotype was found to comprise three distinct subtypes: *MYCN*-amplified, *MYCN* non-amplified high-risk, and *MYCN* non-amplified low-risk signatures^17^. The MES phenotype was significantly enriched in relapsed tumors^17^ and has been linked to chemotherapy resistance^16^.

In this study we applied a pan-cancer, multi-omic approach to elucidate metabolic dependencies in cancer and identified a critical role of DHODH, both as a prognostic marker and as a mediator of tumor cell survival in neuroblastoma. We report that the therapeutic combination of DHODH inhibition and the standard of care chemotherapeutic temozolomide has curative potential in a transgenic neuroblastoma mouse model and is a promising candidate for the treatment of high-risk neuroblastoma.

## RESULTS

### Neuroblastoma cells accumulate nucleotide metabolites and express high levels of *DHODH*

In order to explore metabolic dependencies and characteristics in neuroblastoma, we utilized the Cancer Cell Line Encyclopedia (CCLE) metabolome dataset. Comparing a panel of neuroblastoma cell lines (n = 17) to a wide range of different solid tumor cell lines (n = 746) using partial least-squares discriminant analysis (see Methods), we identified several metabolites that were accumulated in neuroblastoma (**Fig. 1A, Fig. S1A**). These include the pyrimidines 2-deoxycytidine and cytidine, the purine derivative hypoxanthine, and the versatile nutrient glutamine, which provides nitrogen for nucleotide synthesis^18^. Due to the accumulation of 2-deoxycytidine and cytidine, we hypothesized that pyrimidine metabolism may play an important role in neuroblastoma disease and therefore curated a list of 16 genes encoding enzymes related to pyrimidine synthesis and salvage (**Supplementary Table 1**). Next, we studied the vulnerability to CRISPR-mediated genetic perturbation of these enzymes by assessing CERES scores for a panel of 990 cancer cell lines, using the DepMap resource from the Broad Institute (https://depmap.org). A lower CERES score indicates that the cells are more likely to be dependent on that particular gene. The enzymes CAD, DHODH, and UMPS, which catalyze subsequent steps of *de novo* pyrimidine synthesis, were found to correlate strongly to each other in the DepMap dataset, with Pearson correlation scores > 0.7 (**Supplementary Table 1**). Cancer cell lines that are highly dependent on one enzyme of *de novo* pyrimidine synthesis are also highly dependent on the other two, suggesting that such cancers are “addicted” to UMP production and pyrimidines. This strong inter-correlation between CAD, DHODH, and UMPS was also evident in the subset of neuroblastoma cell lines (**Fig. S1B**).

**Figure 1.**
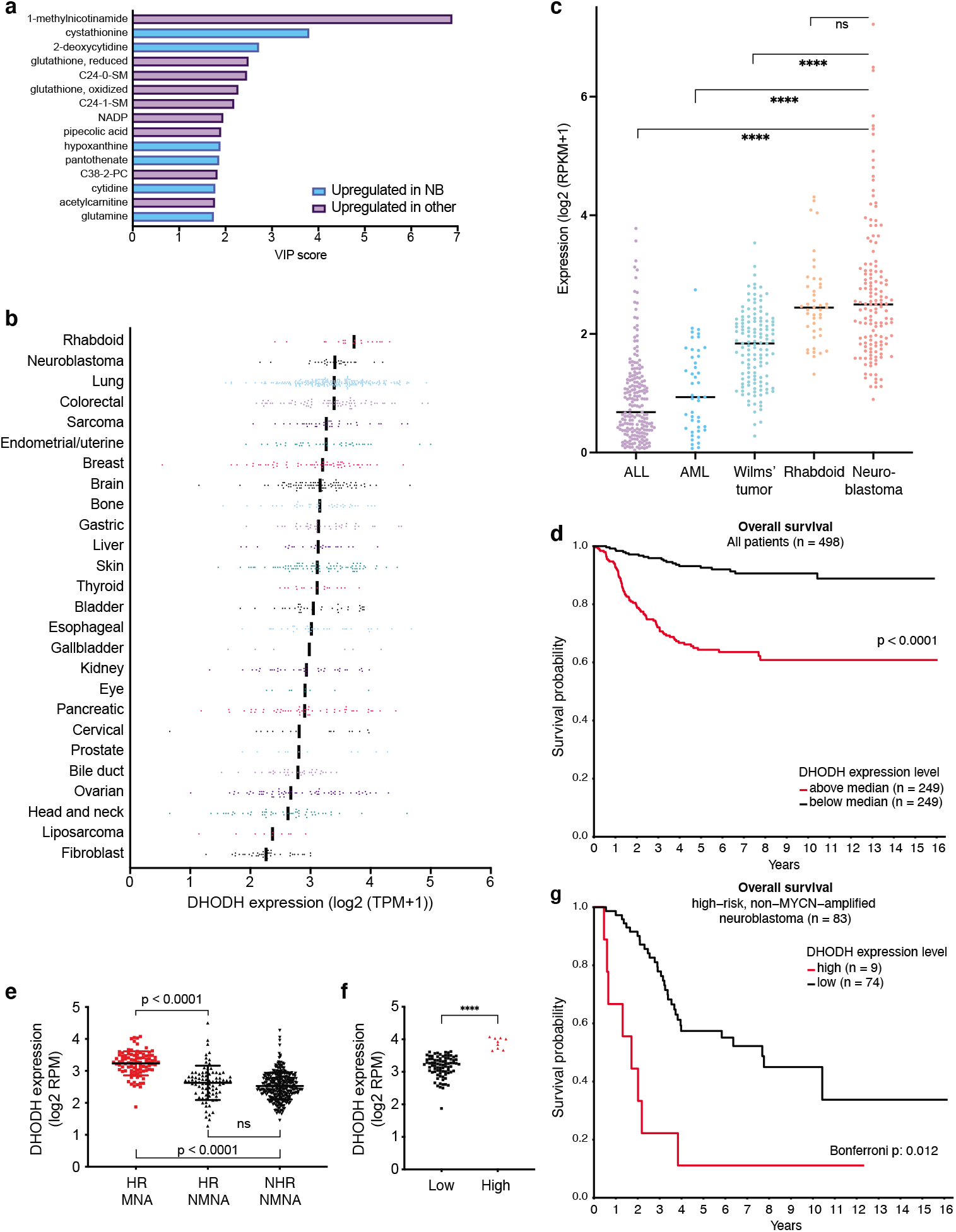
*DHODH* is highly expressed in neuroblastoma and expression correlates with disease stage, prognosis, and survival. (**A**) Top 15 accumulated metabolites in neuroblastoma cell lines compared to other solid tumor cell lines of the CCLE metabolomics dataset. Metabolite enrichment in neuroblastoma is quantified as VIP (variable importance in projection) scores generated through partial least squares discriminant analysis. (**B**) *DHODH* expression in solid tumor cell lines of the CCLE dataset, grouped by primary disease. Black lines represent median values. (**C**) *DHODH* expression in primary pediatric tumor samples of the TARGET dataset. Black lines represent median values. ****: p < 0.0001; ns: not significant (as evaluated by one-way ANOVA analysis with multiple comparisons). ALL: acute lymphatic leukemia; AML: acute myeloid leukemia. (**D**) Overall survival in 498 neuroblastoma patients (SEQC-498 dataset), separated by median *DHODH* expression. Red: *DHODH* above median (high); black: *DHODH* below median (low). (**E**) *DHODH* expression in 498 neuroblastoma samples (SEQC-498 dataset). HR: high-risk, MNA: *MYCN* amplified, NHR: non-high-risk; NMNA: non *MYCN* amplified; ns: not significant. *DHODH* expression is significantly higher in the *MYCN* amplified tumors as evaluated by one-way ANOVA analysis with multiple comparisons. (**F**) *DHODH* expression levels in high-risk, non-*MYCN* amplified tumors. High/Low groups are identified by the KaplanScan feature of the R2 database (http://r2.amc.nl). (**G**) Kaplan-Meier curve showing the survival of the groups in (F). Note that the “highest DHODH group” (n = 9) is associated with a particularly poor overall survival.

In comparison to other solid tumor cell lines (**Fig. 1B**) and primary pediatric tumor samples (**Fig. 1C**), *DHODH* was expressed at high levels in neuroblastoma and rhabdoid tumors. There is increasing evidence that dihydroorotate dehydrogenase (DHODH) is a promising therapeutic target for many different cancer types^19–24^ and DHODH inhibitors are currently under evaluation in several clinical trials. Consequently, we next sought to explore the clinical relevance of DHODH in neuroblastoma.

### DHODH is an independent unfavorable prognostic marker in neuroblastoma

We next evaluated two independent neuroblastoma cohorts to investigate the prognostic implications of *DHODH* expression in human neuroblastoma. High *DHODH* expression was significantly correlated with poor survival when combining all stages of the disease (**Fig. 1D, Fig. S1D**). Interestingly, the highest levels of *DHODH* expression were observed in high-risk MYCN amplified tumors (**Fig. 1E**) and *DHODH* expression increased with INSS stage (**Fig. S1C**). Cox regression analysis of two independent cohorts, with adjustment for age, INSS stage, risk group, and *MYCN* status, confirmed that increasing *DHODH* expression is an independent risk factor significantly correlated with an unfavorable overall survival (**Supplementary table 2**). Notably, even within the high-risk and non-high-risk groups, high *DHODH* expression (above median) correlated significantly with worse survival in non-*MYCN* amplified cases (**Fig. S1F-G**). Importantly, a subset of high-risk, non-*MYCN* amplified cases with very high *DHODH* expression (**Fig. 1F**) demonstrated dismal prognosis; a “highest risk group” (**Fig. 1G**). Taken together, high *DHODH* expression is significantly correlated with adverse outcomes in patients with neuroblastoma.

### DHODH inhibition is an effective therapy in pre-clinical models of neuroblastoma

The effect of DHODH inhibition on tumor cell growth was evaluated in a panel of neuroblastoma cell lines and a non-malignant fibroblast cell line. We used the DHODH inhibitor brequinar, which has demonstrated promising effects in both human and murine AML models^25^ and is currently under investigation in human clinical trials (ClinicalTrials.gov Identifier: NCT03760666). Across a panel of neuroblastoma cell lines brequinar demonstrated suppression of tumor cell growth with IC50 values in the low nanomolar range (**Fig. S2A-B, Supplementary table 3**). One notable exception was the SH-EP cell line, which was highly resistant to brequinar treatment.

To evaluate the therapeutic potential of DHODH inhibition *in vivo*, mice bearing SK-N-BE(2) or SK-N-AS xenografts were treated with brequinar. These cell lines were selected based on their *MYCN* amplification status (SK-N-BE(2): *MYCN* amplified; SK-N-AS: non-*MYCN* amplified) and the difference in sensitivity to brequinar treatment (**Fig. S2A**). Comparable to our *in vitro* observations, brequinar treatment dramatically suppressed tumor growth in SK-N-BE(2) xenografts, with a less pronounced effect in the SK-N-AS xenograft model (**Fig. 2A**).

**Figure 2.**
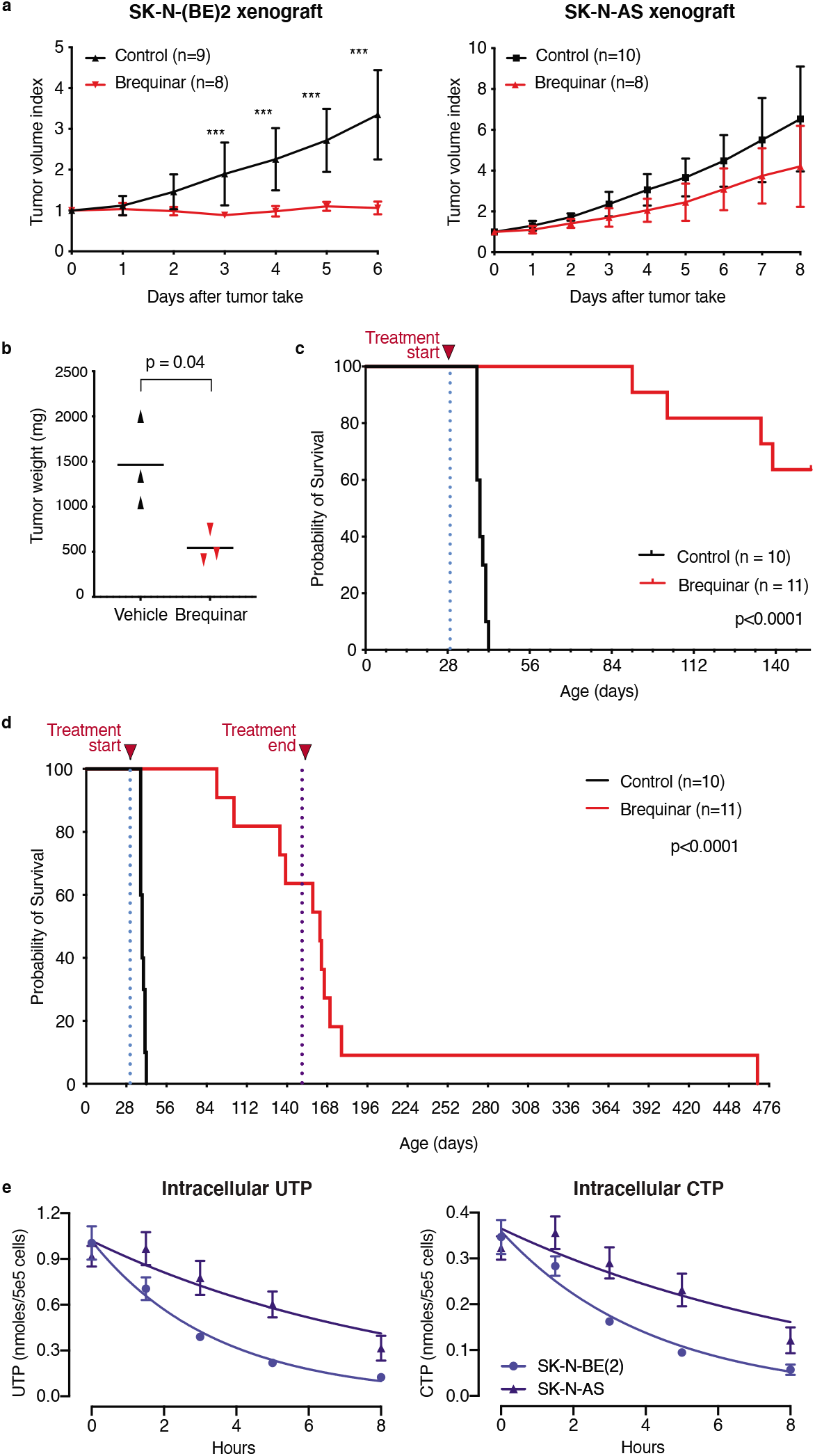
DHODH inhibition reduces neuroblastoma growth *in vitro* and *in vivo*. (**A**) Tumor volumes of SK-N-BE(2) and SK-N-AS NMRI nu/nu xenografts treated with brequinar 50 mg/kg intraperitoneally every 3 days. Tumor volume index at a given day is calculated as the tumor volume relative to the volume at the time of inclusion. Error bars and symbols indicate mean with SD. (**B**) Tumor weights in homozygous TH-MYCN mice 72 hours after receiving one dose of either brequinar or vehicle. Error bars show mean with SD. Groups are compared using unpaired t-test. (**C**) Kaplan-Meier curve of homozygous TH-MYCN mice treated with brequinar 50 mg/kg intraperitoneally. Treatment started at 32 days of age (marked with dotted line) and continued for 120 days when possible. (**D**) Survival of brequinar treated mice after treatment termination (marked with a second dotted line) from the same cohort as shown in (C). (**E**) Inhibition of DHODH decreases the pool of pyrimidines in neuroblastoma cell lines. UTP (left panel) and CTP (right panel) decay in SK-N-BE(2) and SK-N-AS cells following treatment with Brequinar (1 μM).

Since pro-differentiating effects of brequinar have been demonstrated pro-differentiating effects in acute myeloid leukemia^25–27^ we investigated whether the neuroblastoma xenografts still had the potential to relapse following cessation of brequinar therapy. After treatment cessation at day 18, SK-N-BE(2) xenografts quickly recurred, indicating that the malignant cells were not completely eradicated or terminally differentiated (**Fig. S2C**).

To evaluate the therapeutic potential of brequinar in an immunocompetent *in vivo* model, we treated homozygous transgenic TH-MYCN neuroblastoma mice^28^ with brequinar once abdominal tumors were palpable. A single dose resulted in a dramatic reduction of tumor weight as compared to vehicle-treated control mice after just 72 hours (p=0.04, **Fig. 2B**). Extended brequinar treatment in TH-MYCN mice (120 days) led to a strikingly prolonged survival (p<0.0001, **Fig. 2C**). Similar to the SK-N-BE(2) xenografts, tumors eventually relapsed after discontinuation of brequinar treatment (**Fig. 2D**).

### *MYCN* and *DHODH* dependencies correlate in neuroblastoma

Based on our observations *in vivo* (both SK-N-BE(2) cells and the transgenic TH-MYCN mouse model are dependent on *MYCN* expression), we next examined whether *DHODH* dependency correlates with *MYCN* dependency in neuroblastoma. We observed a significant correlation in CERES scores between these genes (**Fig. S2D**), where *MYCN*-dependent cell lines tend to be more dependent on *DHODH*. At 24 hours, brequinar treatment resulted in the downregulation of *DHODH* as well as *MYCN* (SK-N-BE(2)) or *MYC* (SK-N-AS). (**Fig. S2G**). In comparison to SK-N-BE(2), the SK-N-AS cells recovered their *MYC* and *DHODH* expression, suggesting this may be a potential mechanism underlying their relative resistance to DHODH inhibition. A similar phenomenon was observed in brequinar treated xenografts, where *MYCN* (SK-N-BE(2)) or *MYC* (SK-N-AS) expression recovered 72 hours post-treatment (**Fig. S2H**). In the TH-MYCN mouse model, we observed that endogenous *Mycn* was reduced 72 hours after one dose of brequinar. However, expression of the human *MYCN* transgene was not affected after 72 hours (**Fig. S2F**). This may be comparable to the recovery effect of *MYCN* expression seen after 72 hours in SK-N-BE(2) xenografts (**Fig. S2H**).

These results prompted us to hypothesize that *MYCN* expression, and the resulting changes in downstream gene expression and metabolism, might result in an increased dependence on *de novo* pyrimidine synthesis. SK-N-BE(2) and SK-N-AS cells were treated with brequinar and samples were processed for quantitation of intracellular nucleotides by HPLC (**Fig. 2E**). Inhibition of DHODH by brequinar resulted in a more rapid depletion of intracellular nucleotides in the SK-N-BE(2) cells, suggesting an increased dependence on *de novo* synthesis and an inability to adequately salvage extracellular nucleotides to meet cellular demands. Notably, the intracellular purine pools are spared, speaking to the on-target specificity of brequinar (**Fig. S2I**). To control for the potential confounder of cell proliferation rate, a CFSE proliferation assay was performed and demonstrated a highly similar doubling time between the two cell lines (**Fig. S2J**).

### DHODH inhibition preferentially targets the neuroblastoma ADRN cell state

Neuroblastoma cell lines exist in at least two distinct malignant cellular states: MES and ADRN cells^15,16^. In light of this phenomenon, we analyzed gene expression data from CCLE neuroblastoma cell lines and generated ADRN and MES scores using the expression signature scoring as described by van Groningen *et al*^15^. Notably, SK-N-AS shows a distinct MES score compared to other cell lines and we observed a significant inverse correlation between ADRN and MES scores (**Fig. 3A**). A group of six cell lines (SK-N-SH, LS, SK-N-AS, CHP-212, GI-ME-N and KP-N-SI9s) displayed mesenchymal transcriptional profiles with high MES scores and low ADRN scores (**Fig. 3A**). Interestingly, five of these six cell lines had high *DHODH* CERES scores, indicating a lower likelihood of *DHODH* dependency (**Fig. 3B**). Conversely, the NB cell lines that are most dependent on *DHODH* all had low MES scores (**Fig. 3B**). Furthermore, SK-N-AS and SH-EP cells were also the most resistant to treatment with brequinar *in vitro* (**Fig. S2A**) with the sensitivity of SH-EP cells comparable to the non-malignant stromal MRC5 cell line (**Fig. S2A-B**). Of note, SH-EP is known to display a mesenchymal phenotype^15,16^. Moreover, in SK-N-AS cells (high MES score, **Fig. 3A**), we observed a plateau effect where increasing doses of brequinar did not further decrease cell viability (**Fig. S2A**), suggesting that neuroblastoma cells with a predominantly MES phenotype are less dependent on DHODH and more resistant to its inhibition.

**Figure 3.**
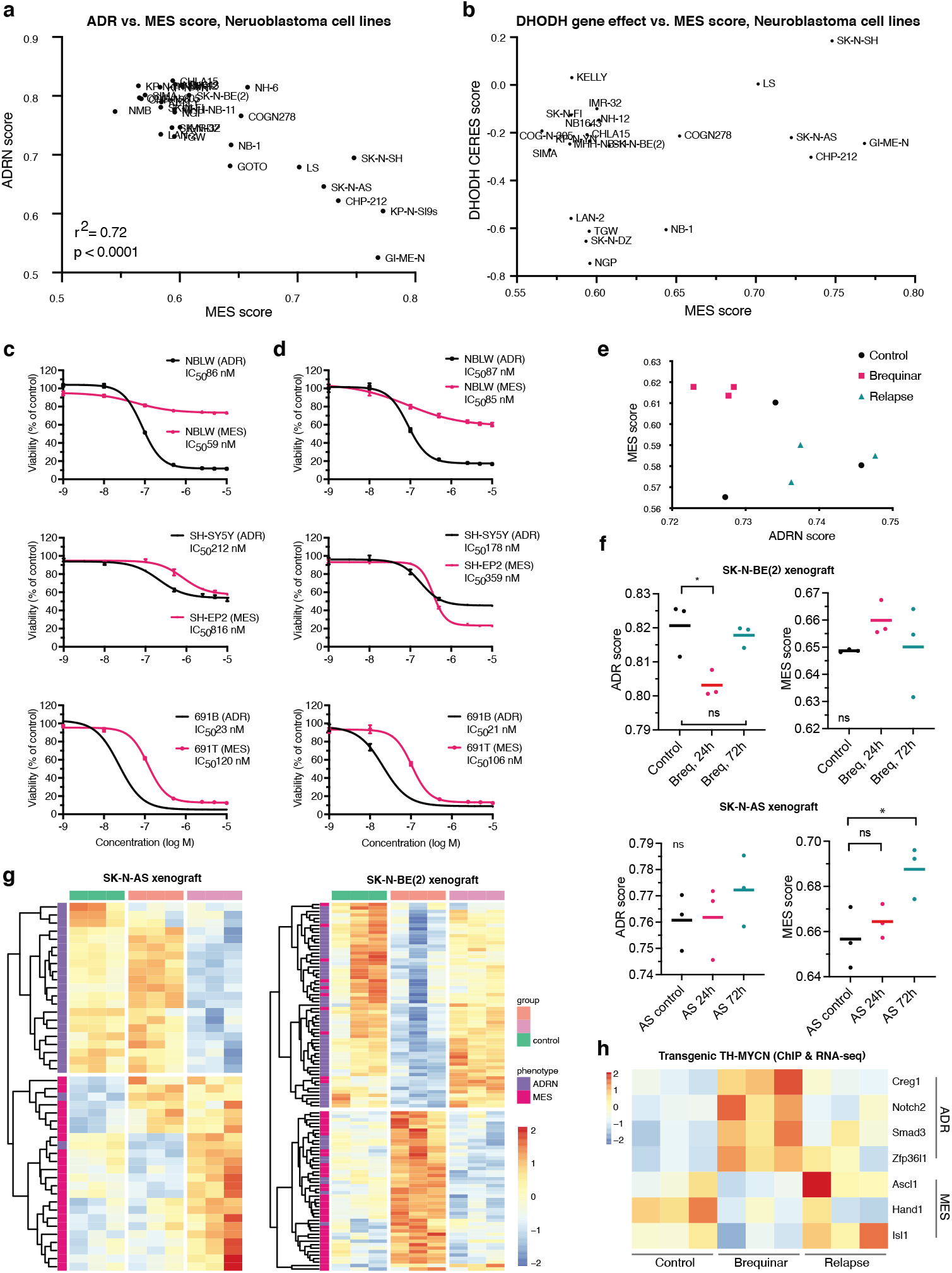
MES cells are more resistant to DHODH inhibition than ADRN cells. (**A**) ADRN and MES scores of neuroblastoma cell lines in the CCLE/DepMap dataset. The two scores are inversely correlated. One distinct group of six cell lines has particularly high MES scores. (**B**) MES gene expression scores and *DHODH* gene dependency scores (CERES) of neuroblastoma cell lines in the CCLE/DepMap dataset. The highest MES-scoring cell lines all display relatively high CERES scores whereas the most *DHODH*-dependent cell lines all display low MES scores: mesenchymal NB cell lines are less dependent on *DHODH*. (**C-D**) MTT viability assay (C) and CyQuant proliferation assay (D) in three isogenic ADRN (black) and MES (pink) neuroblastoma cell line pairs treated with brequinar. The MES counterparts are generally more resistant to brequinar. (**E**) ADRN and MES scores from TH-MYCN tumor samples in three groups: vehicle-treated controls (black), tumors from brequinar-treated mice (sampled 72 hours after one dose; pink), and relapsed tumors after 120 days of brequinar treatment (50 mg/kg every three days; turquoise). Note the shift towards lower ADRN and higher MES scores after brequinar treatment. (**F**) Violin plots of ADRN and MES scores in SK-N-BE(2) and SK-N-AS xenografts sampled 24 and 72 hours after one dose of brequinar (50 mg/kg i.p). In SK-N-BE(2), there is a significant, but transient decrease in ADRN score at 24 hours. In SK-N-AS, the MES score is significantly higher at 72 hours. Groups are compared using one-way ANOVA test with multiple comparisons. *: p < 0.05; ns: not significant. (**G**) Heatmaps showing z scores of significantly deregulated ADRN/MES genes in neuroblastoma xenografts: SK-N-AS 72 hours after one dose of brequinar; SK-N-BE(2) 24 hours after one dose of brequinar. Red color indicates expression higher than average; blue color indicates expression lower than average. In both cases, there is an increased expression of MES genes and a decreased expression of ADRN genes following treatment. (**H**) RNA expression heatmap (z scores) of ADRN- and MES-associated transcription factors in TH-MYCN tumors displaying corresponding changes in H3K27Ac enhancer signal. Red color indicates higher expression than average, blue color indicates lower expression than average. Following brequinar treatment, enhancer signal and gene expression of MES- and ADRN-associated transcription factors increases and decreases, respectively. The pattern at relapse is similar to the pattern in untreated control tumors.

To compare the sensitivity to brequinar within the same cell line model we treated isogenic cell line pairs that reflect ADRN or MES cell states (SH-EP2/SH-SY5Y, 691B/691T and NBLW-ADRN/NBLW-MES^15^). We confirmed that the ADRN neuroblastoma cells were more sensitive to DHODH inhibition in comparison to their MES counterparts (**Fig. 3C-D**), further substantiating our observation that the MES cell state is more resistant to DHODH inhibition.

To explore this phenomenon *in vivo*, we treated homozygous TH-MYCN mice with a single dose of brequinar and harvested tumors after 72 hours. Relapsed tumors (after discontinuation of brequinar treatment) were also sampled for gene expression analysis. 72 hours after one dose of brequinar, we observed a transcriptomic shift towards a more MES-like signature (**Fig. 3E**). Interestingly, at the time of relapse the ADRN/MES expression signatures were comparable to that of untreated controls, suggesting a MES-to-ADRN cellstate shift in relapsed tumors.

We next evaluated the superenhancer landscape, using ChIP-seq to assess H3K27Ac signal) in TH-MYCN tumors at 72 hours after one dose of brequinar. Overall, brequinar-treated and control samples were highly similar in terms of their enhancer profiles (**Fig. S3A-C**) and superenhancers typical of the ADRN neuroblastoma phenotype^15^ were primarily observed in both groups (**Fig. S3C**). To cross-compare ChIP-seq and RNA-seq data, we identified genes where the direction of change in gene expression correlated with the direction of change in H3K27Ac signal in a nearby genomic region (e.g., upregulated genes with corresponding higher levels of H3K27 acetylation). Brequinar treatment resulted in enhancer-related downregulation of genes related to synaptic function and neuron cellular components such as synapses and axons (**Supplementary Table 4**). We also observed enhancer-related upregulation of genes related to TNF signalling and other immune-related processes. Furthermore, seven of the core ADRN/MES transcription factors (TFs) identified by van Groningen *et al*^15^ displayed enhancer-related transcriptional changes. Mesenchymal TFs (*Notch2, Creg1, Smad3, Zfp36l1*) were upregulated after brequinar treatment. ADRN TFs (*Ascl1, Hand1, Isl1*) were downregulated, while the relapse tumors were similar to the controls (**Fig. 3H**).

Gene expression analysis of brequinar treated xenografts at 24h and 72h revealed a similar change in ADRN and MES signatures. In SK-N-AS xenografts, the shift from ADRN towards MES signatures was evident at 72 hours (**Fig. 3F**). In SK-N-BE(2) xenografts, the ADRN scores transiently decreased at 24 hours, but were comparable to controls 72 hours after one dose of brequinar (**Fig. 3F**). Among the ADRN/MES signature genes that were significantly deregulated at 24 (SK-N-BE(2)) or 72 (SK-N-AS) hours, upregulated genes were mainly MES signature genes, whereas downregulated genes were mainly ADRN signature genes in both cell lines (**Fig. 3G**). Taken together, we observed an increased MES and/or decreased ADRN signature after DHODH inhibition *in vivo*. This may be explained by brequinar preferentially inhibiting ADRN tumor cells, shifting the balance between these two populations in the tumor tissue.

### A combination of DHODH inhibition and temozolomide demonstrates curative potential in transgenic neuroblastoma mice

Given that single agent DHODH inhibition was not curative in the xenograft and transgenic *in vivo* models of neuroblastoma, we next sought to combine brequinar with the standard-of-care chemotherapy temozolomide, a DNA alkylating agent, which is a treatment backbone in relapsed and refractory neuroblastoma^29^.

Brequinar and temozolomide demonstrated a synergistic inhibitory effect *in vitro* (**Fig. 4A, S4A-D**). To explore this effect *in vivo* we treated tumor-bearing homozygous TH-MYCN homozygous mice with one dose of brequinar and/or temozolomide and performed tumor immunohistochemistry on tumors labelling proliferating cells (Ki-67) and apoptosis (active caspase 3). At 24 hours, after one dose of either brequinar or temozolomide, we observed a marked decrease in the proportion of Ki67+ cells and an increase in active caspase 3+ cells (**Fig. 4C, S5A-C**). 24 hours after two consecutive days of treatment, first with brequinar, then temozolomide, we observed a strong increase in cells expressing the mesenchymal marker vimentin (**Fig. 4C**).

**Figure 4.**
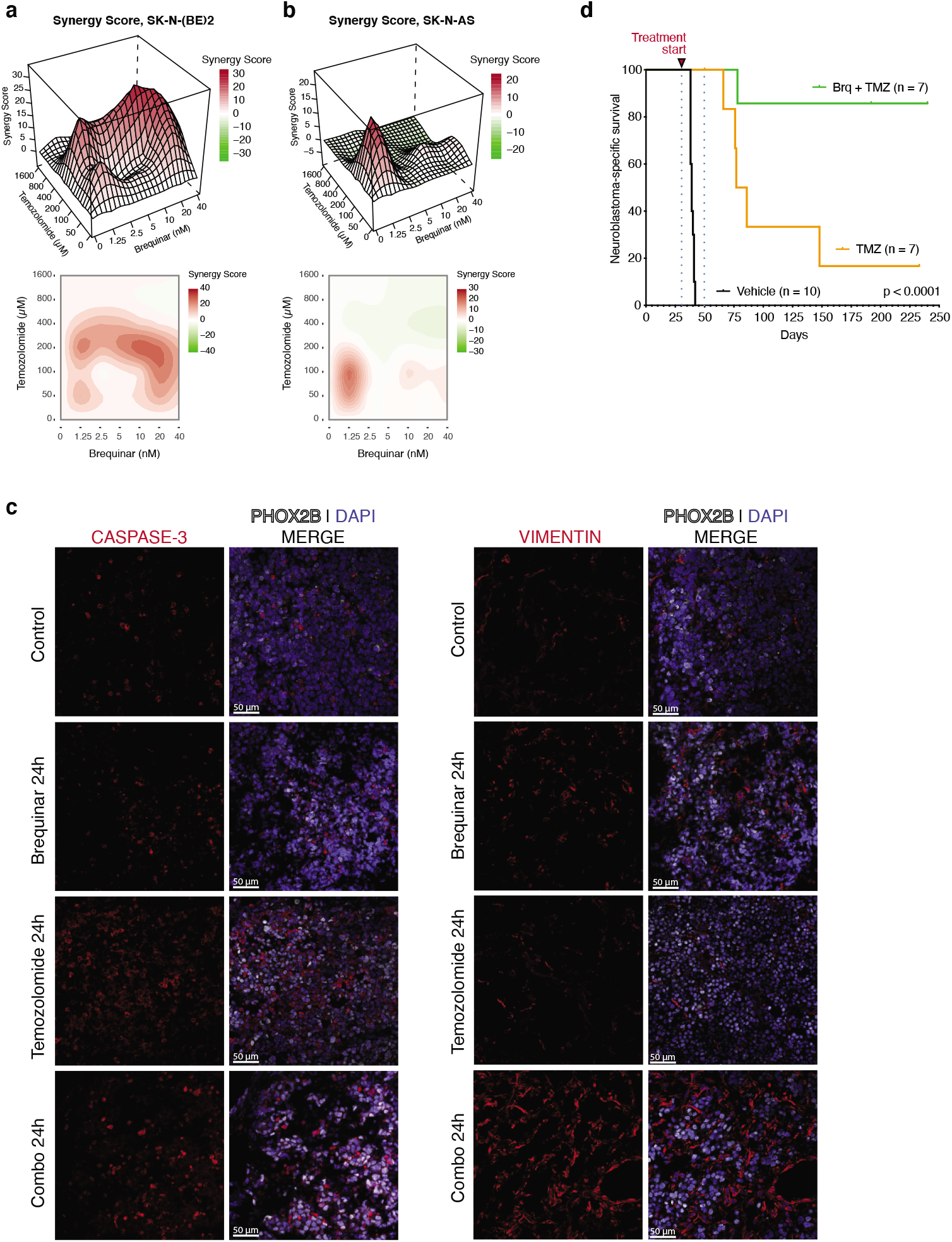
Combinations of brequinar and temozolomide are synergistic in vitro and have curative potential in vivo. (**A-B**) Synergy scores of temozolomide and brequinar combinations in SK-N-AS (A) and SK-N-BE(2) (B) cells treated with both drugs. Synergy scores are based on three independent experiments. Red color indicates synergy. ZIP scores are shown as 3D (upper panel) and 2D (lower panel) representations. ZIP scores indicate the average % excess response due to drug interactions. (**C**) Immunostainings at 40X of active caspase-3 (red), vimentin (red) and PHOX2B (white) in TH-MYCN mice treated with one dose of brequinar and/or temozolomide; sampled 24 hours after the last dose. Nuclei were stained with DAPI (blue). (**D**) Overall survival of homozygous TH-MYCN mice treated with either temozolomide alone (n = 7) or a combination of brequinar and temozolomide (n = 7). Treatment started when mice were 32 days old and was performed according to the overview in Fig. S4E. For comparison, control mice (black) are the same as in Fig. 2C.

To investigate the therapeutic efficacy of this promising combination, we treated homozygous TH-MYCN mice with three doses of brequinar followed by two cycles of temozolomide, or with two cycles of temozolomide alone (**Fig. S4E**). The TH-MYCN mice previously treated with vehicle control were used as a comparator group (**Fig. 2C, 4D**) and these untreated mice generally succumb within approximately six weeks post-birth. When comparing temozolomide- and combination-treated to vehicle-treated mice, survival was significantly prolonged in both groups (p < 0.0001). However, following discontinuation of treatment, all but one of the temozolomide treated mice suffered from relapse. In the combination group, we observed a dramatic prolongation of survival where five out of seven mice did not suffer from neuroblastoma relapse even 190 days after treatment cessation (**Fig. 4D**). Two mice in the combination therapy group ultimately died from thoracic tumors, of which one was a PHOX2B+ thoracic neuroblastoma (suggesting a metastatic or primary thoracic neuroblastoma) and the other a PHOX2B-, N-Myc-, CD45+, CD3+ lymphoma (**Fig. S6**). In conclusion, the combination of brequinar and temozolomide has great curative potential *in vivo*.

## DISCUSSION

Cancer cells activate the purine and pyrimidine biosynthesis pathways to meet the nucleotide demands of rapidly proliferating cells^30^. There is increasing evidence that dihydroorotate dehydrogenase (DHODH), an essential enzyme for de novo pyrimidine synthesis, is a promising therapeutic target in various cancer types, including melanoma, acute myeloid leukemia, glioblastoma, lung cancer, pancreatic cancer and prostate cancer^19–24^. We recently demonstrated that the DHODH inhibitor brequinar is an effective differentiation therapy in myeloid leukemia, demonstrating prolonged survival in mouse and PDX models due to differentiation of leukemic stem cells into mature myeloid cells^25^. Multiple DHODH inhibitors are currently under investigation in human clinical cancer trials (clinicaltrials.gov, identifiers NCT03760666, NCT03709446, NCT02509052). It was recently demonstrated that DHODH inhibition may suppress tumor growth *in vivo* by inducing ferroptosis^31^. However, the mechanisms by which DHODH inhibition induces the differentiation of malignant cells still remain largely unknown.

In patients with high-risk neuroblastoma, there is currently a need for improved therapy, as survival rates remain less than 50%^32^. The aim of this study was to identify novel metabolic vulnerabilities in neuroblastoma and to evaluate the role of DHODH as a potential therapeutic target in this disease. Brequinar was first introduced as an anticancer agent in the 1980s^33^. Several phase II clinical trials were performed in the following decade, evaluating the therapeutic potential of DHODH inhibition in a wide range of adult solid tumor malignancies^34–37^. Although well-tolerated, the clinical efficacy of brequinar was disappointing. Later, the DHODH inhibitor leflunomide was approved as an anti-inflammatory agent in the treatment of rheumatoid arthritis. Interestingly, DHODH inhibition with leflunomide has been shown to block neural crest development and decrease melanoma growth *in vivo*^19^. In addition, it has also been explored as a potential therapeutic agent in the treatment of multiple myeloma^38^. This is of potential interest for neuroblastoma research as both neuroblastoma and melanoma are neural crest-derived cancers^1,39^. Using a large high-throughput chemical screen, we previously identified that DHODH inhibitors were capable of inducing differentiation in acute myeloid leukemia models *in vitro* and *in vivo*^25^.

By analyzing large, publicly available large neuroblastoma patient cohorts, we established *DHODH* expression as an independent prognostic marker in neuroblastoma after adjusting for known adverse factors such as disease stage, age, and *MYCN* amplification. Furthermore, we observed a subset of high-risk, non-*MYCN*-amplified tumors with very high *DHODH* expression levels and particularly dismal prognoses. This indicated a potentially biological role of DHODH in neuroblastoma and prompted us to perform further analyses to evaluate DHODH as a therapeutic target.

In this study, neuroblastoma cells are highly sensitive to DHODH inhibition with IC50 concentrations in the low nanomolar range. Additionally, we observed that the most sensitive cell lines, such as SK-N-BE(2), were the ones with highly ADRN phenotypes^15,16^ and that *MYCN* dependency correlated with *DHODH* dependency. In contrast, cell lines with MES phenotype, such as SK-N-AS and SH-EP, were more resistant to DHODH inhibition. When we compared the *MYCN* amplified cell line SK-N-BE(2) to the non-*MYCN*-amplified SK-N-AS cell line, we observed a greater reduction of pyrimidines UTP and CTP upon brequinar treatment in the SK-N-BE(2) cells. These results may indicate that *MYCN*-driven neuroblastoma cells are less capable of pyrimidine salvage, and more dependent on *de novo* pyrimidine biosynthesis via DHODH. However, even though DHODH inhibition was effective in the ADRN, *MYCN*-driven SK-N-BE(2) cells *in vivo*, monotherapy was not curative and DHODH inhibition did not induce terminal differentiation (**Fig. 2D, S2C**).

The TH-MYCN model is a model of spontaneous and aggressive neuroblastoma that is also driven by *MYCN* overexpression^40^. Although ADRN and/or MES phenotypes have not been previously demonstrated in this model, we found that the superenhancer core regulatory circuitry in these tumors is also largely ADRN, similar to *MYCN* amplified cell lines (**Fig. S3C**). In this model, DHODH inhibition is also highly effective in preventing tumor growth. However, similar to SK-N-BE(2) xenografts, the effect was incomplete and in both model systems, tumors relapsed 1-2 weeks following the discontinuation of treatment.

Brequinar treatment triggered a transcriptomic shift from an ADRN signature towards a MES signature *in vivo*. The MES gene signature after brequinar treatment may represent non-malignant stroma and/or malignant MES neuroblastoma cells and the MES enrichment may be caused by selective killing of ADRN cells, resulting in a higher residual number of stromal and/or mesenchymal neuroblastoma cells. Another possibility is that the MES enrichment is caused by ADRN neuroblastoma cells transitioning towards a more MES phenotype, as has previously been demonstrated *in vitro*^15,41^.

The combination of the alkylating agent temozolomide and the DHODH inhibitor brequinar demonstrated synergistic effects *in vitro* and a striking effect on the survival of homozygous transgenic TH-MYCN mice. Pyrimidine depletion has previously been shown to induce DNA damage and double-strand breaks^42^. Thus, one possible explanation for the observed synergy may be the combined effect of alkylation and pyrimidine depletion on DNA damage and breakage.

In conclusion, our data demonstrate the therapeutic impact of DHODH as a critical mediator of neuroblastoma cell growth. Specific DHODH inhibition combined with the alkylating agent temozolomide is highly effective in neuroblastoma preclinical models. Targeting DHODH as a cancer-specific metabolic dependency may benefit high-risk neuroblastoma patients and is a strong candidate for further testing in clinical studies.

## MATERIALS AND METHODS

### Cancer Cell Line Encyclopedia (CCLE), DepMap, and TARGET gene expression analysis

Clean liquid chromatography-mass spectrometry (LC-MS) metabolomics data of CCLE cell lines was aquired from the publication by Li *et al*^43^. Metabolomics analyses comparing neuroblastoma cell lines to other solid tumor cell lines were performed using the Statistical Analysis module of MetaboAnalyst 5.0^44^. In partial least-squares discriminant analysis, cell lines were grouped as “neuroblastoma” (n = 17) or “other” (n = 746). DepMap/CCLE copy number data (21Q2 Public dataset) was used to define *MYCN* amplification, where cell lines with log2 (relative to ploidy +1) > 3 was considered as amplified. Cell line gene expression (log2 (TPM+1), 21Q2 Public dataset) and CRISPR (DepMap 21Q2 Public+Score, CERES) datasets were acquired via the DepMap portal and annotated according to the primary disease in the corresponding cell line metadata table. Pre-computed statistics for CRISPR data was collected in the DepMap portal. Gene expression values from various primary pediatric cancer samples (log2 (RPM+1), TARGET dataset) were accessed and downloaded through the cBioPortal website (http://www.cbioportal.org). To quantify ADRN and MES scores, gene signatures and the scoring approach outlined by van Groningen *et al*^15^ was used.

### Statistical survival analyses

Survival and gene expression data from two neuroblastoma patient cohorts was collected from the publicly available datasets SEQC-498 (GSE49711) and TARGET-NBL (FPKM-UQ normalized expression data downloaded from portal.gdc.cancer.gov). For the TARGET dataset, corresponding survival data was acquired from a previous publication^45^. Gene expression values were log2-transformed in order to achieve a distribution closer to the Gaussian distribution. The distribution of continuous variables was checked using histograms and normal QQ-plots. Overall survival was defined as days from diagnosis until death from any cause or end of follow-up. Event-free survival was defined as days from diagnosis until disease progression or relapse, death from any cause, or end of follow-up, whichever came first. The association between survival and covariates was evaluated using Cox proportional hazards models. The linearity assumption of continuous covariates was checked by fitting models with restricted cubic splines, with three knots, using the function *rcs* in the R package *rms*. The effect of age at diagnosis was found to be non-linear, and age was thus adjusted for by using restricted cubic splines with three knots, in order to adjust for the whole age effect. All survival analyses were performed using R version 3.5.0.

The ‘‘highest risk” subset of non-MYCN amplified, high-risk neuroblastoma cases (**Fig. 1F-G**) was identified by using the KaplanScan feature of the R2 database (https://r2.amc.nl) with the minimum group size set to 8.

### Cell cultures and reagents

For WST-1 cell viability and qPCR analyses, cell lines were grown in Dulbecco’s modified Eagle’s medium/F12 (Gibco/Thermo Fisher Scientific; SH-SY5Y) or RPMI 1640 (Gibco/Thermo Fisher Scientific; SK-N-AS, SK-N-FI, SK-N-BE(2), SK-N-FI, SK-N-DZ, SK-N-SH, IMR-32, Kelly, SH-EP, MRC5) supplemented with 10% fetal bovine serum, 2mM L-glutamine, 100 μg/ml streptomycin, and 100 IU/ml penicillin G (all from Life Technologies/Thermo Fisher Scientific). Cell lines were purchased from ATCC (ATCC-LGC Standards, Middlesex, UK) and grown at 37 °C in a humidified 5% CO_2_ atmosphere. Cell line identities were verified by short tandem repeat genetic profiling (AmpFISTR Identifiler PCR Amplification Kit; Applied Biosystems), routinely tested for mycoplasma (Mycoplasmacheck, Eurofins Genomics) and used in passages below 25.

For MTT viability and CyQuant proliferation analyses, SH-SY5Y and SH-EP2 cell lines were cultured as previously described^46^. The isogenic cell line pair 691B and 691T was cultured in neural stem cell (NSC) medium as previously described^47^. The NBLW-ADR and NBLW-MES cell lines were derived from the parental NBL-W cell line^48^ through their differential adhesion to culture plates. All NBL-W cell lines were cultured in RPMI-1640 medium supplemented with 10% fetal bovine serum, 1x non-essential amino acids, 20 mM L-glutamine, 10 units/mL penicillin and 10 μg/mL streptomycin (all from Life Technologies / Thermo Fisher Scientific). Cell line identities of isogenic ADRN/MES cell line pairs were verified by short tandem repeat (STR) analysis (Multiplexion). Cell lines were routinely checked for the presence of mycoplasma using the MycoAlert detection kit (Lonza).

### Drugs

Brequinar was a kind gift from Dr. Sykes. Temozolomide was purchased from Sigma-Aldrich. For *in vitro* studies, brequinar and temozolomide were dissolved in DMSO. For *in vivo* studies, brequinar was dissolved in a vehicle consisting of 70% PBS and 30% polyethylene glycol (PEG)-400, pH 7.2, while temozolomide was dissolved in DMSO and further diluted in sterile physiological saline.

### Cell viability and proliferation assays

For WST-1 cell viability analyses, Opti-MEM (Gibco/Thermo Fisher Scientific) was used and supplemented with 1% fetal bovine serum and penicillin, streptomycin, and L-glutamine in the same concentrations as for cell culturing. Cells were seeded in 96 well plates (SK-N-BE(2): 5,000 cells per well, all other cell lines 10000 cells per well). The following day, cells were treated with brequinar and/or temozolomide. After 72 hours, cell viability was evaluated using a colometric formazan-based cell viability assay (WST-1; Roche, Sigma-Aldrich).

Absorbance was measured at 450 and 650 nM using a VersaMax reader (Molecular Devices). Cell viability is presented as % of untreated control cells. All concentrations were at least tested in triplicates and experiments repeated at least three times. IC50 values (inhibitory concentration 50%) were calculated from log concentration-effect curves in Prism (GraphPad Software) using non-linear regression analysis. For temozolomide/brequinar combination experiments *in vitro*, viability concentrations (% of control) with corresponding drug concentrations were analyzed using the *synergyfinder* R package (v2.4.16)^49^.

For CyQuant and MTT assays, SH-SY5Y/SH-EP2 (DMEM; 10000/5000 cells per well), 691B/691T (culturing medium as described above; 15000 cells per well), and NBLW-ADRN/NBLW-MES (RPMI; 15000/7500 cells per well) cell line pairs pairs were seeded in 96-well plates and treated with brequinar in the indicated concentrations. For each drug concentration, sextuplet replicates were used. After 72 hours of treatment, cell DNA content was measured using the CyQuant assay (C35012, Life Technologies/Thermo Fisher Scientific) according to the manufacturer’s instructions with the exception that CyQuant reagents were added at half of the indicated volumes. Metabolic activity was measured by adding 10 μl of 3-(4,5-dimethylthiazol-2-yl)-2,5-diphenyltetrazolium bromide (MTT; 10 mg/ml, Sigma). After four hours of incubation, 100 μl of 10% SDS/0.01M HCl was added to stop the reaction. Optical densities were quantified using a Synergy HT microplate reader (Biotek).

### Incubation of cells with drug for metabolite measurement

SK-N-BE(2) and SK-N-AS cells were incubated with 1 μM of brequinar for different time points. Five hundred thousand cells were harvested for each sample and cell line respectively at the indicated times. After being washed with phosphate buffered saline, cells were processed for nucleotide extraction. Nucleotides were extracted using 0.4M perchloric acid and the extracts were neutralized with 10M KOH and stored at −20 C until analyzed as described previously^50^.

### Measurement of intracellular nucleoside triphosphate by HPLC

The neutralized extracts were applied to an anion exchange partisil - 10 SAX column and eluted at a flow rate of 1.5 ml / min with a 50 min concave gradient (curve #9, Alliance 2695 Separations Module; Waters Corp.) from 60% 0.005 M NH_4_H_2_PO_4_ (pH 2.8) and 40% 0.75 M NH_4_H_2_PO_4_ (pH 3.8) to 100% 0.75 M NH_4_H_2_PO_4_ (pH 3.8). The column eluate was monitored by UV absorption at 262 nm, and the nucleoside triphosphates were quantitated by electronic integration with reference to external standards. The intracellular concentration of nucleotides contained in the extract was calculated from a given number of cells of a determined mean volume. The calculation assumed that the nucleotides were uniformly distributed in a total cell volume.

### CFSE analysis of proliferation rate

Cells were labeled per the manufacturer’s protocol with CellTrace Far Red (Thermo Fisher Scientific). The cells were then assayed daily by flow cytometry to monitor the decay of their geometric mean fluorescence intensity. A viability dye (7-AAD) was included to exclude dead cells. The data were plotted using GraphPad Prism in order to calculate the fluorescence decay and doubling time.

### Quantitative PCR

SK-N-BE (2) and SK-N-AS cells were treated with vehicle or brequinar (180 nM SK-N-AS, 7 nM SK-N-BE(2)) for 24 and 72 hours. Total RNA was prepared from cultured cells using the RNeasy MiniKit (Qiagen) including an on-column DNase treatment with the RNase-Free DNase Set (Qiagen). Isolated RNA was transcribed into cDNA with the High-Capacity RNA-to-cDNA Kit (Applied Biosystems). For amplification, the C100 Touch (Bio-Rad) thermocycler was used. mRNA level was assessed using the TaqMan Universal PCR Master Mix (Applied Biosystems) with the sequence-specific probes DHODH (Hs00361406_m1), MYCN (Hs00232074_m1), HPRT1 (Hs02800695_m1) and SDHA (Hs00188166_m1). Relative expression was calculated according to the delta-delta Ct Method normalized to housekeeping genes HPRT1 and SDHA. For *MYCN*, mRNA expression level was normalized to expression in the MRC5 fibroblast cell line. For relative *DHODH* expression, mRNA levels were normalized to untreated controls.

### Animal studies

Female NMRI nu/nu mice of four to six weeks of age were injected with 5 million SK-N-AS or SK-N-BE(2) cells subcutaneously in the flank and monitored daily for tumor growth. Tumors were measured by caliper and tumor volume was calculated using the formula (width)^2^ × length × 0.44. For each tumor, tumor volume index (TVI) was defined as tumor volume relative to inclusion volume. TVI was calculated each day until the first control was sacrificed. At tumor take, when tumor volume exceeded 200 mm^3^, mice were block randomized into two groups (treatment or control). Mice in the treatment group received 50 mg/kg brequinar every third day by intraperitoneal injection for a maximum of 7 doses. The animals were monitored daily for signs of toxicity including weight loss. Mice were sacrificed when the tumor reached a total volume of 2000 mm^3^ or the tumor exceeded length or width of 200 mm. After sacrifice, tumors were dissected in smaller parts, and either fresh frozen, fixed in 4% paraformaldehyde or stored in RNAlater (Invitrogen).

Transgenic TH-MYCN mice were kept on the 129X1/SvJ background^40^. Genotypes were determined from ear tissue biopsies using qPCR with specific probes designed for wild-type and the *MYCN* transgene (Transnetyx, Cordova, TN, USA). In the first treatment study (**Fig. 2E**), homozygous mice at 32 days of age were block randomized into two groups which received intraperitoneal injections of either vehicle (70% PBS, 30% PEG-400) (n = 10) or brequinar (50 mg/kg) every third day (n = 11) for 120 consecutive days. In the second experiment (**Fig. 4D**), at 32 days of age, homozygous mice were block randomized into two groups: the combination group (n = 7) or the temozolomide group (n = 7). Mice in the combination group were treated with brequinar every third day (50 mg/kg) for a total of 3 doses (treatment day 0, day 3, day 6) followed by 20 mg/kg temozolomide intraperitoneally for a total of 10 doses (treatment days 7-11 and 14-18). Mice in the temozolomide group were treated with temozolomide intraperitoneally (20 mg/kg) at treatment days 0-4 and 7-11 for a total of 10 doses. For timepoint sampling of homozygous TH-MYCN tumors subject to single-agent treatment, mice with palpable abdominal tumors received one intraperitoneal dose of either temozolomide (20 mg/kg), brequinar (50 mg/kg), or vehicle (70% PBS, 30% PEG-400). Mice were then sacrificed and tumors were sampled after 6, 24, 48, or 72 hours. For sampling of homozygous TH-MYCN tumors subject to combination treatment, mice with palpable abdominal tumors received one intraperitoneal dose of brequinar (50 mg/kg) on day 0, one intraperitoneal dose of temozolomide (20 mg/kg) on day 1, and mice were sacrificed on day 2.

All animals were maintained at a maximum of six animals per cage and given sterile water and food ad libitum. The animal experiments were approved by the Stockholm ethics committee for animal research (reference numbers N231/14, N42/14, 5163-2019 and 13820-2019), appointed and under the control of the Swedish Board of Agriculture and the Swedish Court. The animal experiments presented herein were in accordance with national regulations (SFS no. 2018:1192 and SFS no. 2019:66).

### Sectioning and immunofluorescent staining of murine neuroblastoma tumors

Tumors from TH-MYCN mice were dissected and cut to smaller pieces (3 x 3 x 3 mm). The tumor pieces were washed in ice cold PBS and fixed in 4% paraformaldehyde in PBS (pH 7.4) at 4°C for 1 hour. Fixed tumor pieces were washed in PBS and cryoprotected in 30% sucrose in PBS at 4°C for 3 hours with gentle agitation. Subsequently, tumor samples were embedded in OCT and frozen at −80°C. Each tumor sample was sectioned (12 μm) using LEICA CM3050S (CT −20°C, OT −15°C), collected on SuperFrost Plus Adhesion microscope slides (Thermo Fisher Scientific), and stored at −20°C.

For immunostaining, the slides with tumor sections were removed from −20°C and brought to room temperature to dry for 30 min. Heat induced target epitope retrieval was performed by heating 1x Target Retrieval Solution (Dako) to 99°C, submerging slides into near-boiling target retrieval solution, then allowing for it to cool down to room temperature. Slides were washed in PBS containing 0,1% Tween-20 (PBS-T) 3 times, for 10 min each. Primary antibodies diluted in PBS-T were added to the sections and incubated overnight at 4°C in a dark, humid chamber. Sections were then washed in PBS-T 3 times, for 10 min each; followed by incubation with secondary antibodies with 4’,6-diamidino-2-phenylindole (DAPI) diluted in PBS-T for 90 min at room temperature, in a dark, humid chamber. Finally, sections were washed in PBS-T 3 times, for 10 minutes each and then mounted using Mowiol mounting medium (Merck). The primary antibodies used were: PHOX2B (R&D Systems AF4940; 1:40), KI67 (Invitrogen MA5-14520; 1:250), active caspase-3 (R&D Systems AF835; 1:50), and Vimentin (CST 5741; 1:100). For primary antibody detection, secondary antibodies were used, raised in donkey against goat or rabbit conjugated with Alexa-488, or - 555 fluorophores (Invitrogen A32814, A32794) in 1:1000 dilutions. Immunostaining images were acquired using LSM700 confocal microscope (Zeiss) with 40x (water) and 63x (oil) objectives. Images were acquired in .lsm format and Imaris software was used for further analysis.

### Immunohistochemistry

Following resection, tumor tissue was fixed with formaldehyde (3.7-4.0% w/v, AppliChem) for 24h. Fixed tissues were processed in an automated tissue processor (LOGOS, Milestone) and embedded in paraffin. FFPE sections, 4 μm thick, were mounted on glass slides (Superfrost+, Thermo Scientific) and heated for 3 hours at 56 °C. Following deparaffinization in Neo-Clear^®^ (Sigma-Aldrich, Merck) for 2×15 min and rehydration in series of graded alcohols, HIER was performed in Target Retrieval Solution, Citrate pH 6.1 (Dako, Agilent) using a Decloaking Chamber (Biocare Medical) set to 5 min at 110 °C. Unspecific antibody binding sites were blocked with 5% goat serum (Sigma-Aldrich) in TBS containing 0.1% Tween 20 (TBST) (Sigma-Aldrich) for 1h at room temperature (RT). After one wash in TBST, endogenous peroxidase activity was quenched by incubation with BLOXALL^®^ (Vector Laboratories) for 10 min at RT followed by 2×5 min washes in TBST. Tissues were incubated with primary antibodies (PHOX2B: 1:1000, ab183741, abcam; n-Myc: 1:500, 51705 Cell Signaling Technology; CD45: 1:1000, 70257 Cell Signaling Technology; CD3: 1:2000 A0452 Dako/Agilent) diluted in TBST+5% goat serum in a humid chamber overnight at 4°C. Following 3 x 5min washes in TBST, the slides were incubated in ImmPRESS^®^ HRP Goat Anti-Rabbit IgG Polymer Detection Kit (Vector Laboratories) for 30 min at RT. This was followed by 3 x 5min washes in TBST. For visualization, slides were incubated with ImmPACT^®^ DAB Substrate (Vector Laboratories) for 3-5 min. The sections were counterstained with hematoxylin (abcam) followed by dehydration with graded alcohols and Neo-Clear^®^. Slides were mounted using SignalStain^®^ Mounting Medium (Cell Signaling Technology). Sections were scanned at 40x using the Axio Scan.Z1 Digital Slide Scanner, ZEISS. Image analysis was performed using QuPath-0.2.3^51^.

### ChIP and RNA sequencing

Mouse tumor tissue used for RNA sequencing was stored in RNAlater solution (Thermo Fisher, Waltham, MA, USA) at −80 °C, whereas tissue for ChIP sequencing was snap frozen and stored in liquid nitrogen. For brequinar-treated TH-MYCN tumors, tumor tissue RNA/DNA extraction, library preparation and RNA/ChIP sequencing was performed by Active Motif (Carlsbad, CA, USA). Briefly, total RNA was isolated using the RNeasy Mini Kit (Qiagen). RNA quality was assessed using BioAnalyzer 2100 and/or TapeStation RNA Screen Tape (Agilent) and Qubit fluorometric assay (Thermo Fisher Scientific), with RIN values ranging from 6.8 to 9.4. 42 bp paired-end libraries were prepared using the TruSeq stranded protocol and sequenced on the NextSeq 500 platform (Illumina) for a depth of 44.1 – 56.1 M read pairs.

ChIP experiments and library preparation was performed by Active Motif using the HistonePath workflow. Mouse tumor tissues were fixed with 1% formaldehyde for 15 min and quenched with 0.125 M glycine. Chromatin was then isolated by adding lysis buffer followed by tissue disruption with a Dounce homogenizer. Lysates were sonicated and DHA sheared to an average length of 300-500 bp. Genomic DNA (input) was prepared by treating aliquots of chromatin with RNase, proteinase K and heat for de-crosslinking, followed by ethanol precipitation. DNA was quantified on a NanoDrop spectrophotometer. 30 μg chromatin was precleared with protein A agarose beads (Invitrogen) and genomic DNA regions of interest isolated using 4 μg of H3K27Ac antibody (Active Motif, cat# 39133). Complexes were washed, eluted with SDS buffer, and subjected to RNase and proteinase K treatment. Crosslinks were reversed by incubation overnight at 65 °C, and ChIP DNA was purified by phenol-chloroform extraction and ethanol precipitation. Quantitative PCR reactions were carried out in triplicates on specific genomic regions using SYBR Green Supermix (Bio-Rad). Illumina sequencing libraries were prepared from ChIP and input DNA by end-polishing, dA-addition and adaptor ligation. After PCR amplification, DNA libraries were sequenced on the NextSeq 500 platform (Illumina), 75 bp reads, single-end.

Total RNA was extracted from SK-N-AS and SK-N-BE(2) xenograft tissue using the RNeasy Mini Kit (Qiagen) and quality control was performed with Agilent Tapestation according to the manufacturer’s instructions. Libraries were prepared using the TruSeq Stranded mRNA protocol. Library QC was performed using Qubit (Thermo Fisher Scientific) and Agilent Tapestation platform. Indexed libraries were sequenced on the NextSeq 500 platform (Illumina), generating 75bp single-end reads. Basecalling and demultiplexing was performed using CASAVA software with default settings.

### Bioinformatic analyses: ChIP- and integrated ChIP/RNA-seq; TH-MYCN tumors

ChIP- and combined ChIP/RNA-seq analysis of TH-MYCN tumor samples were performed by Active Motif. For enhancer analyses, 75 bp reads were mapped to the mouse genome using the BWA algorithm with default settings. Duplicate reads were removed and only uniquely mapped reads with mapping quality >= 25 were used for further analyses. Using Active motif software, alignments at 3’ ends were extended *in silico* to a length of 200 bp which was the average genomic fragment length of the size-selected libraries. Extended alignments were assigned to 32-nt bins across the genome and the number of fragments per bin was determined. H3K27Ac peaks were determined using the MACS algorithm (v2.1.0)^52^ with p value cutoff 1e-7. Peaks in the ENCODE blacklist of known false CHiP-seq peaks were filtered out. Tag numbers of all samples within a comparison group (i.e controls or brequinar-treated samples) were reduced by random sampling to the number of tags present in the smallest sample. To identify super enhancers, a proprietary Active Motif algorithm was used. MACS peaks were merged if their inner distance was <= 12500 bp. The 5% merged peak regions with the strongest signals were identified as super enhancers. For gene annotation, genes within 25 kb of the super enhancer regions were included.

For the integrated RNA-seq and ChIP-seq analysis of TH-MYCN tumors, 42 bp paired-end RNA-seq reads were aligned to the genome using the STAR algorithm^53^ with defaulted settings. Gene expression was quantified by counting reads aligned to individual genes: only read pairs with both ends aligned to the same strand of the same chromosome were counted. Gene annotations were obtained from the Subread software package^54^ (v1.5.2). Raw read counts were analyzed using DESeq2^55^ for normalization, filtering, and differential calling. ChIP-seq data was reanalyzed using DESeq2^55^ adapted for ChIP-seq data, which yielded normalized read counts, fold-change and p values for all merged regions, thereby comparing H3K27Ac signal intensity between brequinar-treated samples and controls. Merged regions were annotated with genes located within a 10 kb distance. The output was then cross-compared with the RNAseq results. Genomic regions with both ChIP-seq and RNA-seq adjusted p values < 0.1 and shrunken log2 fold change (absolute value) > 0.3 were kept for further analysis. Genes where H3K27Ac and gene expression signal had different directions (i.e log2 fold change had different signs) were filtered out. Finally, for functional enrichment analysis, genes were ranked according their RNAseq padj values and analyzed using gProfiler^56^ (ordered query) and the GO:BP/GO:CC gene sets from MSigDB (version 7.4) and KEGG pathways (2021-05-03 release).

### Bioinformatic analyses: RNA-seq; xenografts and TH-MYCN tumors

Raw sequencing .fastq files from xenografts were aligned to the GRCh38 reference genome; .fastq files from TH-MYCN tumors were aligned to a combined mm10/hg19 genome (10x Genomics, release 3.0.0). Raw reads were aligned using the STAR 2-pass approach^53^. Aligned reads were quantified using htseq-count^57^. Differential gene expression analyses were performed using DESeq2^55^. Genes were ranked according to the adjusted p value and the sign of the log fold change, and gene set enrichment analyses were performed using GSEA^58^. To quantify ADRN and MES scores, gene signatures and the scoring approach outlined by van Groningen *et al*^15^ was used.

## Acknowledgments

We would like to thank the core facility for Bioinformatics and Expression Analysis, which is supported by the board of research at the Karolinska Institutet and the research committee at the Karolinska University Hospital. DBS was supported by the Massachusetts General Hospital Transformative Scholars Program. Thank you to dr. Varsha Gandhi and Mary Ayres at the MD Anderson Cancer Center for HPLC measurement of intracellular nucleotides.

---

**Supplementary figure 1.**
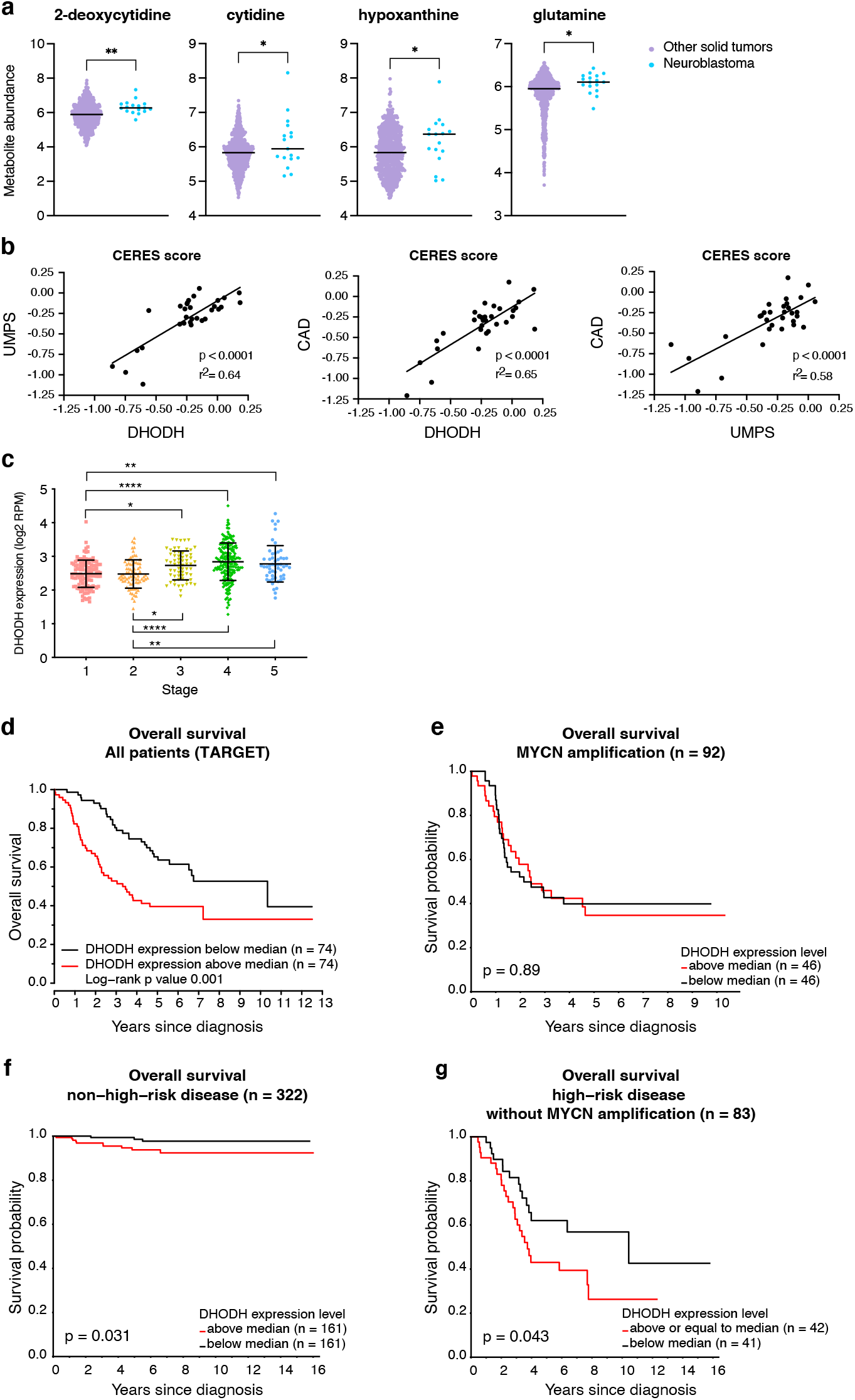
DHODH is highly expressed in neuroblastoma and related to stage, prognosis, and survival. (**A**) Concentrations of nucleotide metabolites significantly accumulated in neuroblastoma compared to other solid tumor cell lines (CCLE metabolomics dataset). Groups are compared using unpaired t-test. *: p < 0.05; **: p < 0.01. (**B**) Correlation between *UMPS, CAD*, and *DHODH* dependency in neuroblastoma cell lines, DepMap/CCLE dataset. Correlation analyses are performed using simple non-linear regression. Dependency on the three essential enzymes of *de novo* pyrimidine biosynthesis is highly inter-correlated. (**C**) DHODH expression in SEQC-498 dataset, separated by INSS stage. Groups are compared using one-way ANOVA with multiple comparisons. *: p < 0.05, **: p < 0.01, ****: p < 0.0001. Lines and error bars indicate mean with SD. (**D**) Kaplan-Meier curve of overall survival in high DHODH vs low DHODH group (separated by median), TARGET dataset. (**E-G**) Kaplan-Meier curves corresponding to the groups in Main Fig 1E showing overall survival separated by DHODH levels above (red) or below (black) median. High DHODH levels are associated with significantly poorer prognosis in the high-risk, non-MYCN amplified and in the non-high risk groups, but not in the *MYCN* amplified group.

**Supplementary figure 2.**
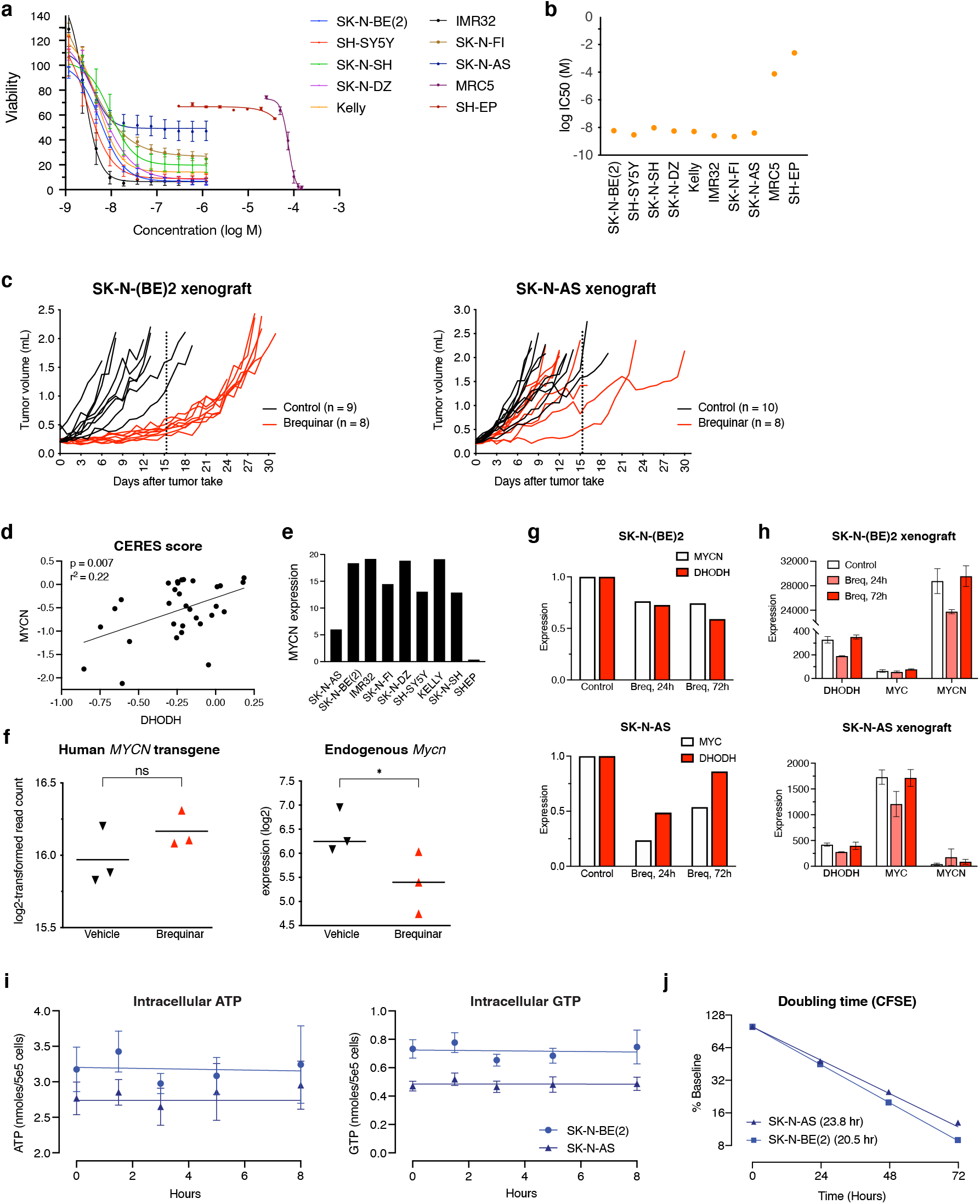
DHODH inhibition reduces neuroblastoma growth *in vitro* and *in vivo*. (**A**) Dose-response curves of WST-1 assays in brequinar-treated cell lines (72h timepoint). (**B**) log IC50 values calculated from non-linear regression of the data shown in (A). (**C**) Tumor volumes of SK-N-BE(2) and SK-N-AS xenografts treated with brequinar (red). Dotted line indicates cessation of treatment. Each line represents one mouse. (**D**) Scatter plot demonstrating correlation between *MYCN* and *DHODH* gene dependency in neuroblastoma cell lines (DepMap/CCLE dataset). (**E**) Expression of *MYCN* in neuroblastoma cell lines, relative to the non-malignant MRC5 fibroblast cell line, as measured by qPCR. (**F**) Expression of human (transgene) *MYCN* and endogenous murine *Mycn* in TH-MYCN tumors 72 hours after one dose of brequinar, measured by RNA-seq. Lines indicate mean values. ns: not significant; *: p < 0.05, as evaluated by unpaired t test. (**G**) Expression of *DHODH* and *MYCN* genes in SK-N-AS and SK-N-BE(2) cell lines treated with brequinar for 72 hours, measured by qPCR. Expression levels are shown relative to untreated controls. (**H**) Expression of *DHODH, MYC* and *MYCN* in SK-N-AS and SK-N-BE(2) mouse xenografts 24 and 72 hours after one dose of brequinar. Expression levels are shown as normalized read counts (RNA-seq). For both cell lines, there is a transient drop in *DHODH* expression at 24 hours. In SK-N-BE(2), there is a transient drop in *MYCN* expression at 24 hours, whereas there is a transient *MYC* drop in SK-N-AS xenografts at 24 hours. (**I**) Inhibition of DHODH does not affect the pool of purines in neuroblastoma cell lines. ATP (left panel) and GTP (right panel) levels in SK-N-BE(2) and SK-N-AS cells following treatment with brequinar (1 μM). (**J**) CFSE assay demonstrating highly similar doubling time in SK-N-AS and SK-N-BE(2) cells.

**Supplementary figure 3.**
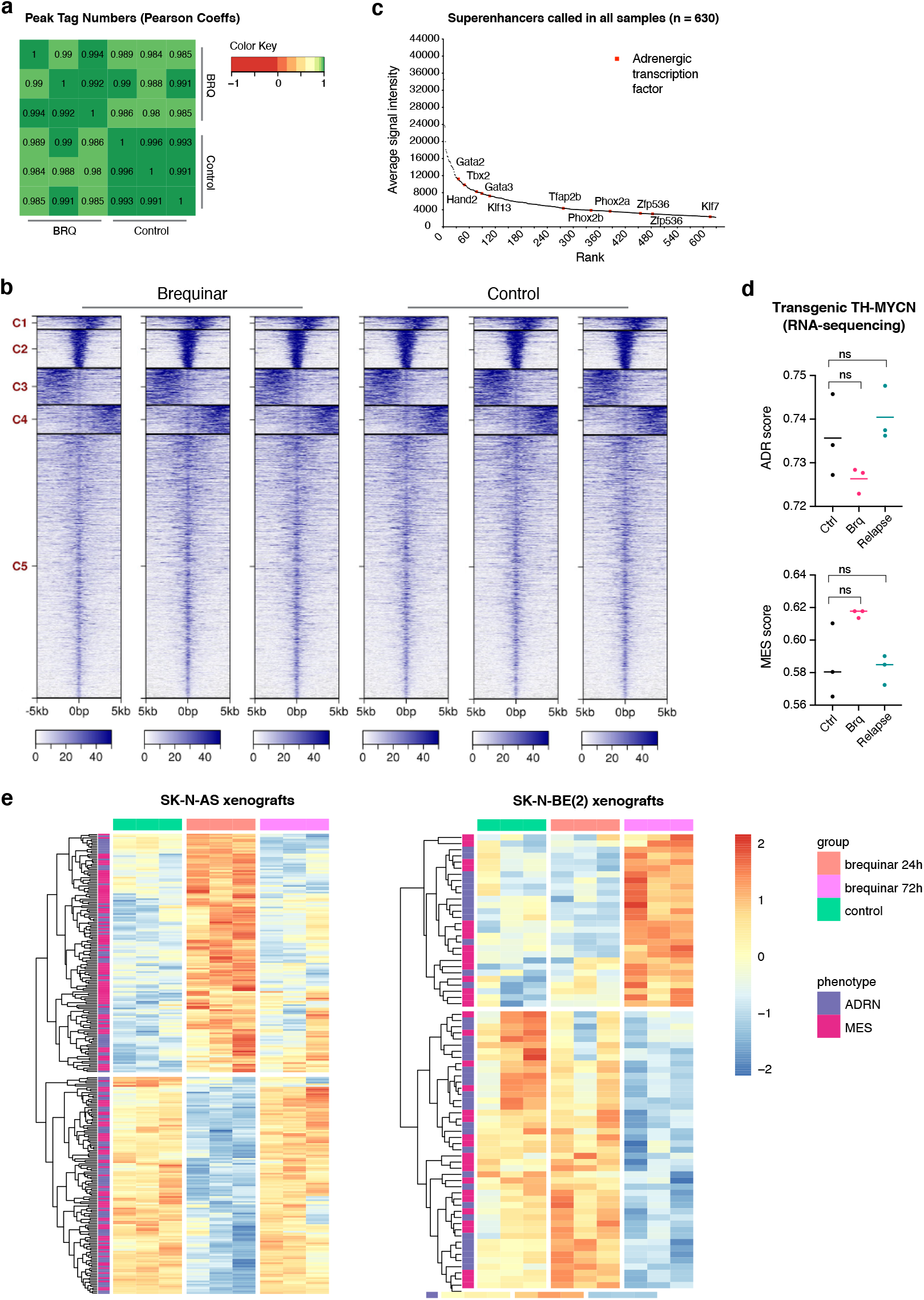
DHODH inhibition induces a shift in MES/ADR gene expression profiles *in vivo*. (**A**) ChIP-seq data of ThMYCN tumors 72 hours after one dose of brequinar. Heatmap showing Pearson correlations between H3K27Ac peak diatributions in all sample-wise comparisons. Although the greatest similarities (darker color) are seen within replicates, overall, all samples are highly similar with coefficients > 0,98. (**B**) Heatmaps showing 5 clusters (C1-C5) of merged peak regions across all samples. Color indicates intensity of H3K27 signal (blue = higher signal). Each blue line along the y axis identifies one particular genomic peak region. X axis indicates the distribution of signal across the region: 0 = the center of the region. (**C**) Superenhancer ranking of 630 SEs detected in all six samples (three controls and three brequinar-treated). Core adrenergic TFs are highlighted. (**D**) ADRN and MES scores in TH-MYCN tumors 72 hours after one dose of brequinar (red) and in relapsed tumors after 120 days of brequinar treatment (turquoise). (**E**) Significantly deregulated ADRN/MES genes 24 hours after one dose of brequinar in SK-N-AS xenografts and 72 hours in SK-N-BE(2) xenografts. Heatmap shows z scores of RNA-seq gene expression data. Red color indicates gene expression above average; blue color indicates expression below average. There is no clear pattern with regards to up- or downregulation of ADRN/MES signature genes.

**Supplementary figure 4.**
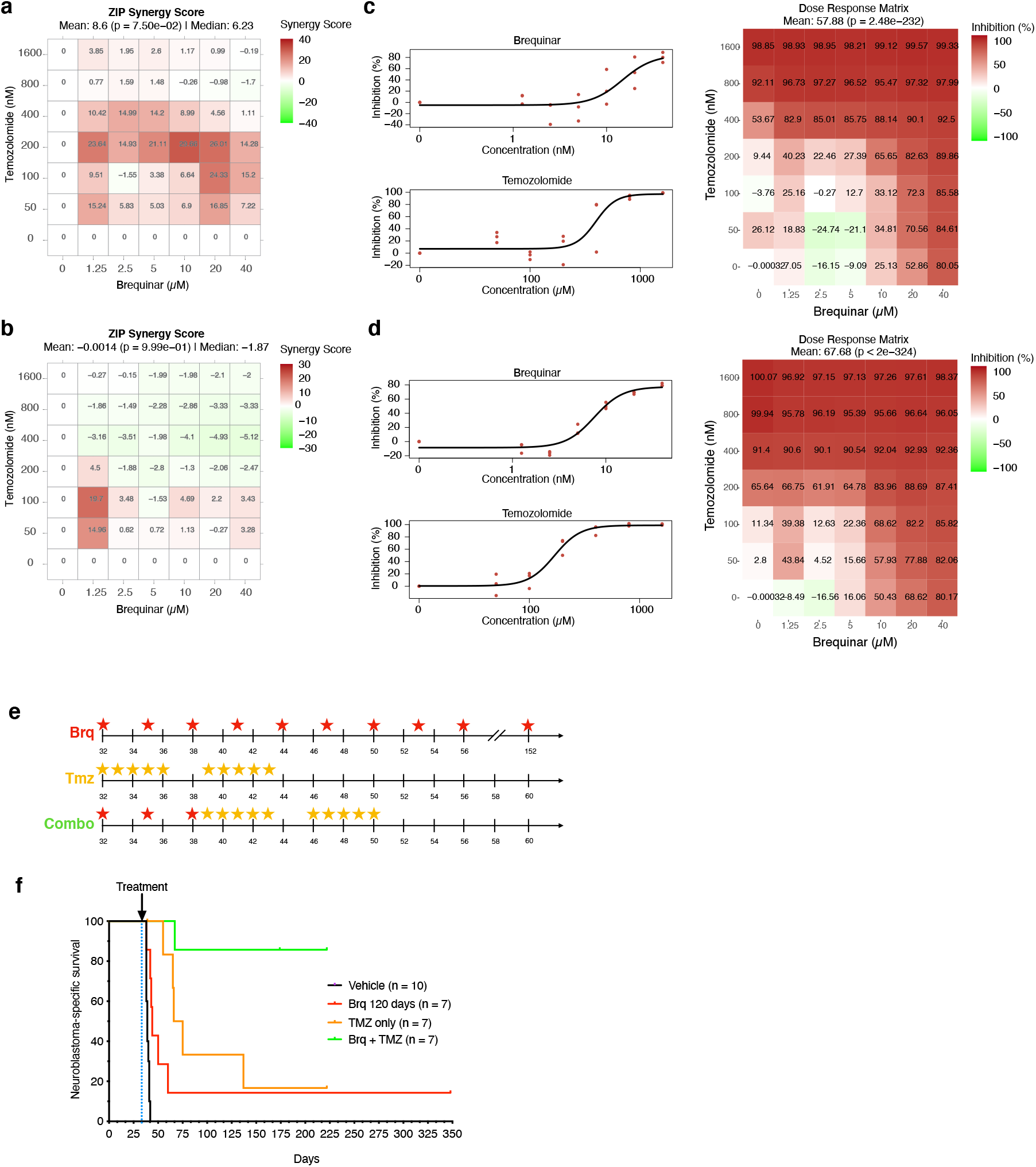
Temozolomide and brequinar display synergistic effects *in vitro* and *in vivo*. (**A-B**) *In vitro* synergy plots from WST-1 cell viability assays in SK-N-BE(2) (panel A) and SK-N-AS(2) (panel B) cells treated with a combination of temozolomide and brequinar. Each cell of the heatmap represents one value in the combination matrix. ZIP scores, quantified by SynergyFinder analysis, are based on three independent replicate experiments. 3D representation of the same data is shown in fig. 4A-B. Brequinar concentrations are shown on the x axis; temozolomide concentrations on the y axis. Red color intensity indicates synergistic effect. (**C, D**) Dose-response curves and heatmaps from the brequinar/temozolomide *in vitro* drug combination experiment. SK-N-BE(2): panel C, SK-N-AS: panel D. (**E**) Schematic overview of brequinar (Brq) and temozolomide (Tmz) treatment regimen in transgenic TH-MYCN mice. Days after birth is shown on the x axis. Stars indicate treatment days. Brequinar dose: 50 mg/kg i.p; temozolomide dose: 20 mg/kg i.p. (**F**) Combined Kaplan Meier curve showing survival of transgenic TH-MYCN mice treated with brequinar and/or temozolomide. Vehicle- and brequinar-treated mice are also shown in Fig. 2D; TMZ- and combination mice are also shown in Fig. 4C.

**Supplementary figure 5.**
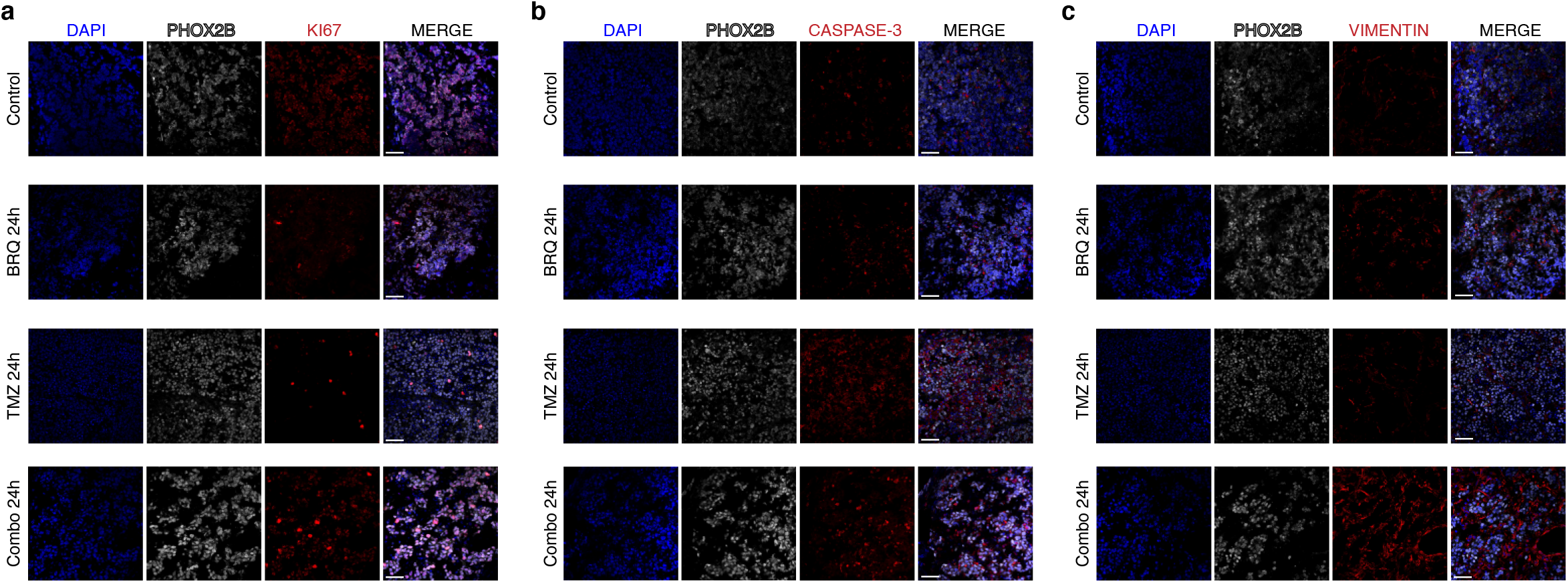
(**A-C**) Immunostainings (40X) of active caspase-3 (red), Ki-67 (red), vimentin (red), and PHOX2B (white) in tumors from TH-MYCN mice treated with one dose of brequinar and/or temozolomide, sampled 24 hours after the last dose. Nuclei were stained with DAPI (blue). Scale bars indicate 50 um.

**Supplementary figure 6.**
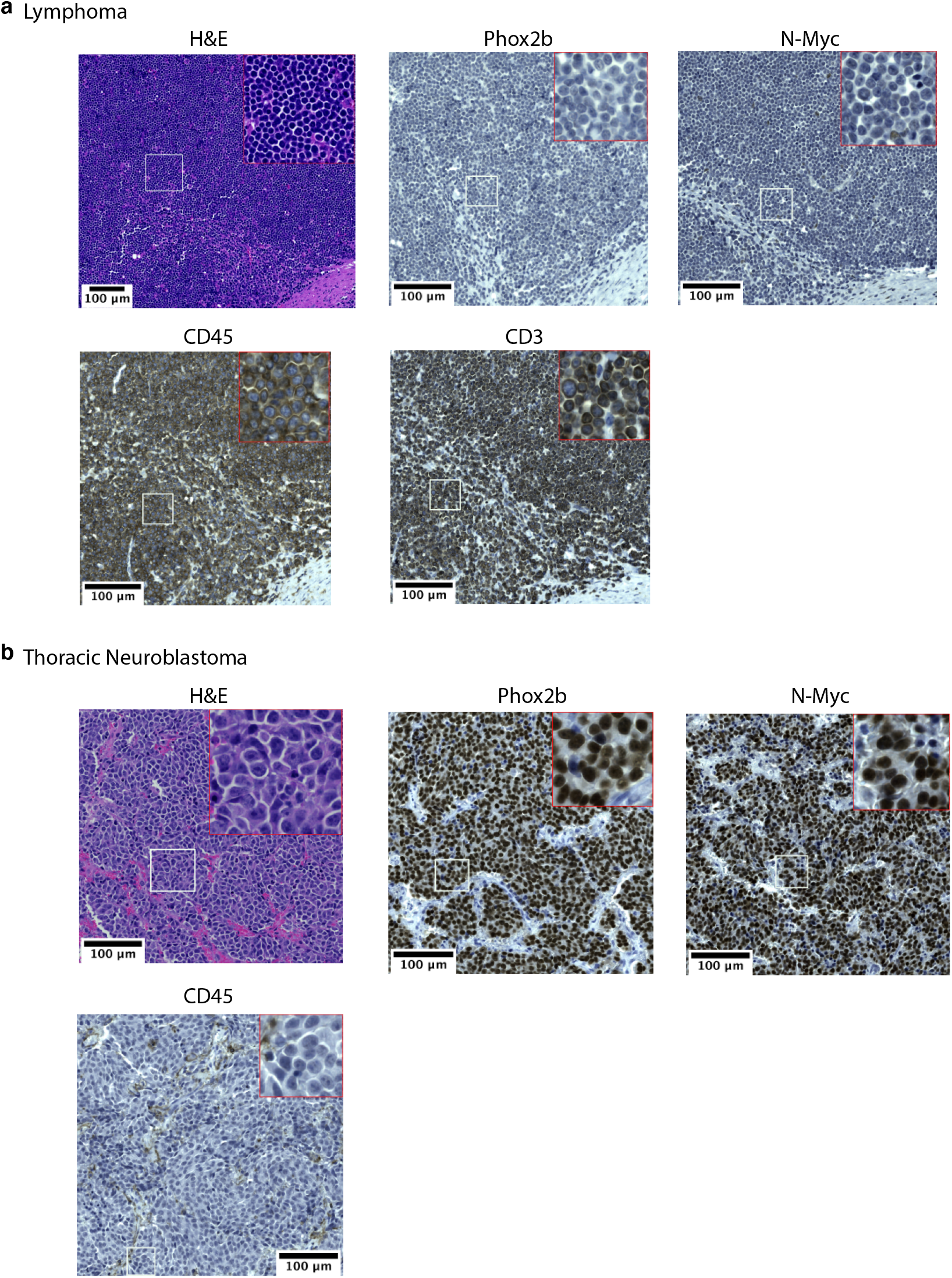
Immunohistochemistry staining of two thoracic tumors seen in TH-MYCN mice treated with three doses of brequinar followed by two courses of temozolomide. (**A**): Thoracic lymphoma negative for PHOX2B and MYCN, strongly positive for CD45 and CD3. (**B**): Thoracic neuroblastoma strongly positive for PHOX2B and N-Myc with some scattered CD45+ immune cells.

**Supplementary Table 1.** Curated list of genes encoding key enzymes of pyrimidine *de novo* synthesis and salvage. For each gene, corresponding gene dependency data in a total of 990 cancer cell lines (DepMap 21Q2 dataset) is provided. For each enzyme that is listed as a dependency in at least one cell line, the top 5 co-dependencies with Pearson correlates are also provided.

**Supplementary Table 2.** Results from Cox regression analysis of *DHODH* expression in the SEQC-498 and TARGET datasets.

**Supplementary Table 3.** IC50 values and *MYCN* amplification status in a panel of neuroblastoma cell lines.

**Supplementary Table 4.** Combined ChIP-seq and RNA-seq analyses of brequinar-treated TH-MYCN tumors. Modified output from DESeq2 analysis. Listed genes display significant gene expression changes with corresponding H3K27Ac signal changes. gProfiler sheets list functional gene enrichment analysis of up- and downregulated genes, respectively.

**Supplementary Table 5.** RNA-seq analysis of brequinar-treated mouse xenografts. Significantly deregulated genes for each timepoint is provided. Corresponding gProfiler functional gene enrichment analyses are added to each gene list.

